# Antigenic evolution on global scale reveals potential natural selection of SARS-CoV-2 by pre-existing cross-reactive T cell immunity

**DOI:** 10.1101/2020.06.16.154591

**Authors:** Chengdong Zhang, Xuanxuan Jin, Xianyang Chen, Qibin Leng, Tianyi Qiu

**Affiliations:** Shanghai Public Health Clinical Center, Fudan University, Shanghai 201508, China; Affiliated Cancer Hospital & Institute of Guangzhou Medical University, State Key Laboratory of Respiratory Diseases, Guangzhou Medical University, Guangzhou, Guangdong 510095, China

## Abstract

The mutation pattern of severe acute respiratory syndrome-coronavirus 2 (SARS-CoV-2) is constantly changing with the places of transmission, but the reason remains to be revealed. Here, we presented the study that comprehensively analyzed the potential selective pressure of immune system restriction, which can drive mutations in circulating SARS-CoV-2 isolates. The results showed that the most common mutation sites of SARS-CoV-2 proteins were located on the non-structural protein ORF1ab and the structural protein Spike. Further analysis revealed mutations in cross-reactive epitopes between SARS-CoV-2 and seasonal coronavirus may help SARS-CoV-2 to escape cellular immunity under the long-term and large-scale community transmission. Meanwhile, the mutations on Spike protein may enhance the ability of SARS-CoV-2 to enter the host cells and escape the recognition of B-cell immunity. This study will increase the understanding of the evolutionary direction and warn about the potential immune escape ability of SARS-CoV-2, which may provide important guidance for the potential vaccine design.

## Introduction

The coronavirus disease 2019 (COVID-19) epidemic caused by a novel Severe Acute Respiratory Syndrome Coronavirus 2 (SARS-CoV-2) was stated in late December 2019 and now become worldwide pandemic^1,2^. The whole-genome sequence of SARS-CoV-2 was first released on Jan 2020^3,4^ and followed by massive strain isolates derived from human patients shared in Global Initiative on Sharing All Influence Data (GISAID)^5^. Coronaviruses (CoVs) are naturally host in vertebrate including birds, swine, cat, multiple bats and human^6^. Based on our best knowledge, there are 7 human-susceptible CoVs and among them, HCoV-229E, HCoV-HKU1, HCoV-NL63 and HCoV-OC43 could only cause slight respiratory infections^7^. On the contrary, other three CoVs including MERS-CoV, SARS-CoV and SARS-CoV-2 could cause serious clinical symptoms of fever, cough, dyspnea, and pneumonia, further, may results in progressive respiratory failure and even death^3^’^8^. Previous research indicated that SARS-CoV-2 share only 50% identity with the MERS-CoV and 79% identity with the SARS-CoV^1,9^, but share over 90% similarity with a bat SARS-related coronavirus (SARSr-CoV, RaTG13) collected in Yunnan, China^3,10^.

The rapid accumulation of the whole-genome sequence^1,5^ illustrated that the mutations are already observed in both structure and non-structure proteins including ORF1ab, envelope (E), Membrane (M), Nucleocapsid (N), Spike (S), ORF3a, ORF6, ORF7a, and ORF7b on SARS-CoV-2. Even though the isolates share with over 99% identity^2^, the mutations could lead to not only the occurrence of different subtypes^2^, but also impact the pathogenicity of SARS-CoV-2^11^. However, the driver of the mutations is still un-revealed and comprehensive investigation is needed to reveal the evolutionary pressure and virulent of the future types of SARS-CoV-2.

The evolution of viruses was mainly affected by the population-level of epidemics, which could allow the opportunities for viruses under evolutionary selective pressure by the immune system of its host^12^ For example, the new antigenic variants could be observed in every 3 to 5 years for influenza A/H3N2 viruses under the selective pressure of the antibody mediated humoral immunity^12^. On the other hand, as an essential part of human immune system, cellular immunity could response to the viral infectious through T-cell and human leukocyte antigen (HLA)^13^. Unlike humoral immunity which is majorly targeting the main antigen protein the viruses, the HLA could recognize the T-cell epitope of the whole genome including structure and non-structure proteins. Moreover, HLA could present the T-cell epitope to the surface of the antigen presenting cell (APC), and further, binding with the T-cell receptors (TCRs) to trigger the immune response^13^. The long-term symbiosis of virus and its host will lead to HLA mediated cellular immunity and accelerate the evolution of virus in multiple cases such as human immunodeficiency virus (HIV) and hepatitis D virus (HDV)^14,15^. Although SARS-CoV-2 is a new emerging pathogen, seasonal HCoVs had been circulating in community for a long time. Currently, multiple works suggesting the existence of cross-reactive T-cell recognitions between circulating seasonal HCoVs and SARS-CoV-2^16-18^. Thus, the long-term community transmission of seasonal HCoVs may also drove the rapid evolution of SARS-CoV-2 under the selective pressure of the immune system. In this work, a comprehensive analysis of the SARS-CoV-2 mutants and the potential binding affinity of T-cell epitopes were provided based on current released mutation strains. Especially for the mutations on ORF1ab which could alter the binding affinity of cross-reactive peptides between seasonal HCoVs and SARS-CoV-2, and the mutations on S protein which could enhance the binding affinity of ACE2 and decrease the binding affinity of antibody. The study presented in this work could not only provides a new insight into the factors driving the evolution of SARS-CoV-2 and its evolutionary pattern, but also provide theoretical guidance for the vaccine and therapeutic antibody design.

## Results

### Workflow of mutation pattern analysis for SARS-CoV-2

Till now, the worldwide community spreading of SARS-CoV-2 could lead to the monitoring and responding of human immune system including HLA mediated cellular immunity and B-cell receptor (BCR) mediated humoral immunity. Consistently, to escape the acquired immune system, the virus will be mutated under the selective pressure. For cellular immunity, both HLA-I and HLA-II molecules presenting multiple alleles in different ethnicities. The diversity of alleles will lead to the presentation of different peptides. Thus, the HLA mediated selective pressure is closely related to the spreading among various human races. By taking benefit from the worldwide spreading of SARS-CoV-2, it is a great opportunity for us to reveal the immune system mediated selective pressure and the potential evolutionary direction of SARS-CoV-2.

Here, we provided a comprehensive analysis in four levels including: 1) mapping the mutations on the whole genome sequence of SARS-CoV-2 and deriving all potential T-cell epitopes (PTEs) involving mutation sites (**Figure 1a**), 2) analyzing the potential peptides based on the circulating regions of viruses worldwide and the local dominant alleles (**Figure 1b**), 3) revealing the selective pressure of HLA through cross-reactive peptides (CRPs) between seasonal HCoVs and SARS-CoV-2 (**Figure 1c**), and 4) evaluating the binding affinity of S protein mutants against human ACE2 and binding antibody (**Figure 1d**). Results indicated that: 1) the mutations were occurred in the whole genome of SARS-CoV-2 including all structure and non-structure proteins, the most frequent mutation sites are in the structure protein of Spike (S) and the non-structure protein of ORF1ab, 2) the frequent mutation sites and strains were discovered in countries such as the United States, the United Kingdom, which have suffered from long-term and large-scale community transmissions, 3) the CRPs between seasonal HCoVs and SARS-CoV-2 on ORF1ab trend to mutate towards the direction of escape from immune response by reducing the binding affinity against both HLA-I and HLA-II, and 4) the mutations of Spike protein could enhance the ability to enter the host cells by increase the binding affinity of ACE2, moreover, escape the surveillance of humoral immunity by decrease the binding affinity of binding antibody.

**Figure 1.**
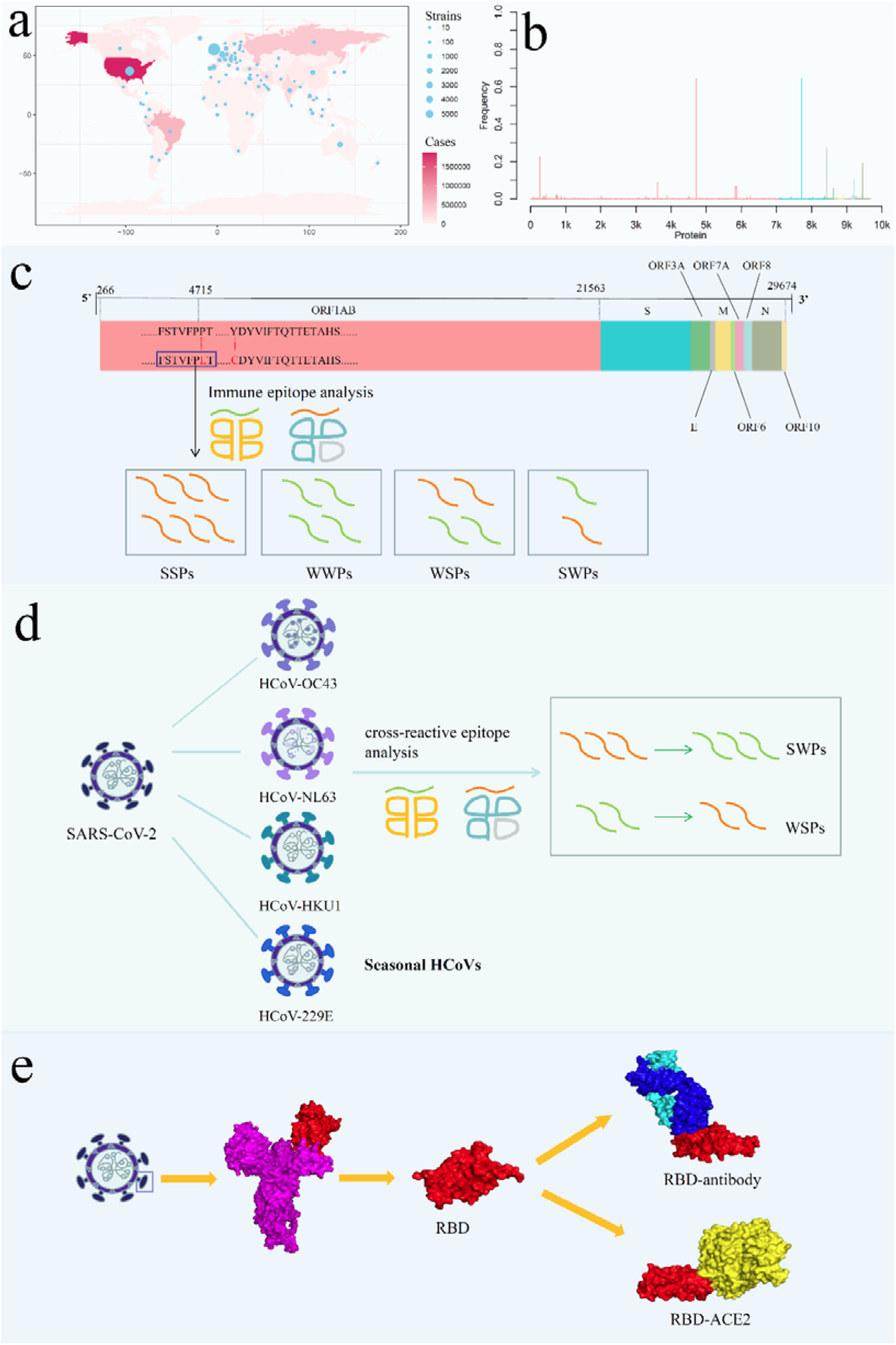
Analysis the mutation pattern of SARS-CoV-2. (a) Collecting mutation strains of SARS-CoV-2 in worldwide scale. (b) Mapping the mutations on the whole genome sequence of SARS-CoV-2. (c) Deriving potential T-cell epitope involving mutations. (d) Revealing the selective pressure of cross-reactive epitopes between seasonal HCoVs and SARS-CoV-2 according to the circulating regions of viruses and the local dominant alleles. (e) Evaluating the binding affinity of S protein mutants against human ACE2 and binding antibody CR3022.

### Frequent mutations on ORF1ab and S protein of SARS-CoV-2

Till May, 20^th^, 2020, we observed 4,420 mutation sites on 10 structure and non-structure proteins including ORF1ab, Spike (S), ORF3a, Envelope (E), Membrane (M), ORF6, ORF7a, ORF8, Nucleocapsid (N) and ORF10 (**Figure 2a**). The conserved regions of all above proteins could be determined by the Identify Conserved Domains (ICD) tools in NCBI^19^ Results showed that ORF1ab, S, ORF3a, M, ORF7a, ORF8 and N contain both conserved and non-conserved area. Among them, ORF1ab and S protein contains the most frequent mutation sites with mutation frequency over 0.6 (**Figure 2a**), which indicating the potential immune pressure mediated evolution. The mutation frequency and counts of each residues could be found in **Supplementary Table 1**. It can be noted that ORF1ab is the largest non-structural polyprotein of SARS-CoV-2, which could trigger the cellular immunity by divided peptides presented by HLA molecules. S protein is the major protein target in humoral immunity, which is mutated under the pressure of BCR or antibodies.

**Figure 2.**
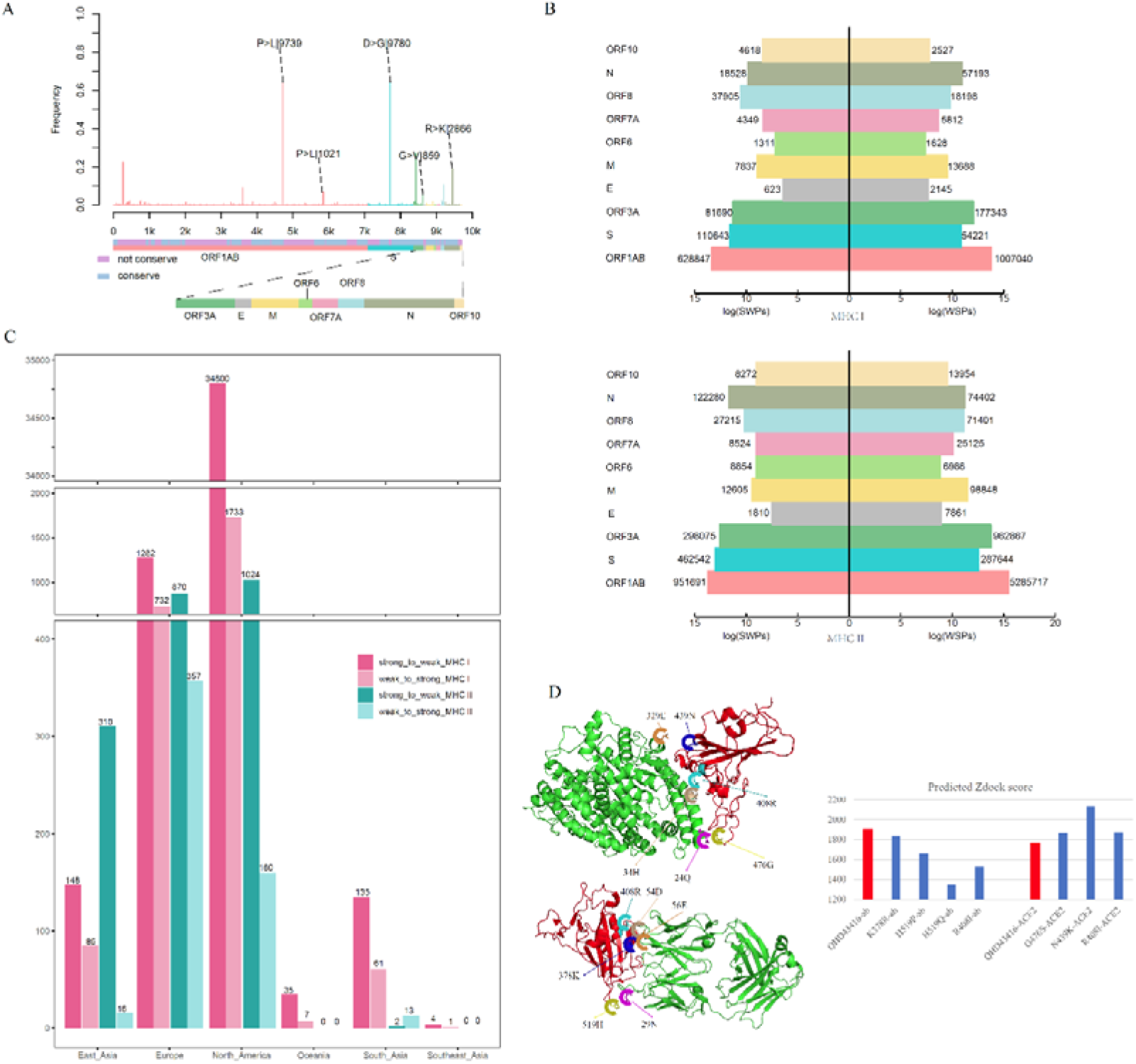
Mutation profile and potential selective pressure analysis of SARS-CoV-2. (a) Mutation patterns on the whole genome of SARS-CoV-2. (b) Number of SWPs and WSPs on the whole genome of SARS-CoV-2. (c) Number of cross-reactive SWPs and WSPs on ORF1ab. (d) Impact of binding affinity for mutations on the S protein.

Further, we counted the continent-specific mutation sites and found the most frequent mutations occurred on S protein, ORF1ab, N protein and ORF3a with 9 sites mutated over 200 times, 5 sites mutated over 800 times and 2 sites mutated over 9,000 times. Among top 5 mutations, D614G on S protein is counted for 9,780 times, followed by P4715L on ORF1ab (9,745 times), R203K on N protein (2,866 times), P5828L on ORF1ab (1,021 times) and G251V on ORF3a (859 times). Detailed information of all mutation sites can be found in **Supplementary Table 2**. More interestingly, the most frequently mutated sites were detected from strains circulating in Northern America and the Europe, which are two major epidemic regions with long-time and large-population transmission of SARS-CoV-2. For example, D614G mutations on S protein were counted for 2,950 times and 3,071 times in the United States and the United Kingdom, respectively. As well as the P4715L mutations on ORF1ab, which were counted for 2,958 and 3,070 times in the United States and the United Kingdom, respectively (**Supplementary Table 3**). The frequent mutations on S protein and ORF1ab might be caused by the selective pressure of humoral immunity and cellular immunity after the long-term, large-scale community transmission of COVID-19.

### Mutations shifting the immunogenicity of T-cell peptides

With the worldwide spreading of COVID-19 epidemic, the infectious time and infected nations or regions are expended. Here, all 8-mer to 15-mer peptides with mutation sites were derived from circulating isolates (***see Methods***). For each strain with collected locations, the HLA alleles with the dominant population coverage (allele frequency ≥ 0.05) were selected and the binding affinity of its corresponding HLA-I/HLA-II alleles were predicted by IEDB standalone MHC-I/MHC-II binding prediction tools^20,21^ (***see Methods***). Reference peptides on sequence of SARS-CoV-2 strain Wuhan-Hu-1 (MN908947.3) was used for comparison (***see Methods***).

Results illustrated that most of the mutations on potential T-cell epitopes (PTEs) didn’t shift: the predicted binding affinity (**Supplementary Table 4 & Supplementary Table 5**). For mutations which alter the binding affinity, weak to strong peptides (WSPs) are significantly higher than strong to weak peptides (SWPs). As been illustrated in **Figure 2a**, the number of WSPs are higher than SWPs for both HLA-I and HLA-II PTEs on most of the non-structure and structure proteins contains. On the contrary, for S protein, the number of SWPs are 2.2 times and 1.8 times higher than WSPs for both HLA-I and HLA-II PTEs, respectively.

### HLA mediated immune pressure could promote the evolution of cross-reactive epitopes between seasonal CoVs and SARS-CoV-2

Moreover, we derived the cross-reactive epitopes (CREs) between seasonal human coronavirus (HCoVs) and SARS-CoV-2 for further analysis (***see Methods***). The infection of seasonal HCoVs are ubiquitous and could cause slight symptoms such as common cold^7,22,23^. Epidemiological data indicating an average of two episodes of “common cold” per year in the adult population, which extrapolated that HCoVs could infected adults in every two to three years^24^. The consistent transmission of HCoVs in human population could maintain the immune memory and produce selective pressure for the CREs between HCoVs and SARS-CoV-2. Here, we detected the CREs between HCoVs and SARS-CoV-2, which might already be circulated across the community before the epidemic of COVID-19. By matching the information of original infectious regions with the dominant HLA-I/HLA-II alleles, 197/358 alleles and 9327/943 peptides which covering 13/11 nations in 7/4 continents were derived for further analysis. For each CREs, the binding affinity of HLA-I/HLA-II with the corresponding alleles were predicted by IEBD MHC-I/MHC-II binding prediction tools (*see **Methods***).

It can be found that, the number of WSPs is 1.6 times than SWPs on ORF1ab for all peptides involving mutations (**Supplementary Table 6**). However, the number of CREs illustrated opposite results, in which cross-reactive SWPs are significantly higher than those of WSPs on all 6 continents including North America, Europe, South Asia, East Asia, Southeast Asia, and Oceania (**Figure 2c**). For example, the results in North America showed that cross-reactive HLA-I SWPs (34,800) are 20.1 times than WSPs (1,733). Similar results could also be found for Europe, South Asia, East Asia, Southeast Asia, and Oceania, in which the number of cross-reactive HLA-I SWPs are 1.75 to 4 times than those WSPs. The SWPs with counting number of 10 can be found in **Supplementary Table 7**.

For HLA-II, CREs could only be detected in North American, Europe, East Asia, and South Asia, which illustrated the same phenomenon. Here, 2,206 cross-reactive HLA-II SWPs were detected, 4 times than those WSPs (546). For North America, Europe, and East Asia, the number of cross-reactive HLA-II SWPs are 2.4 times to 19.4 times than those WSPs. On the contrary, we detected 13 cross-reactive HLA-II WSPs in the data of South Asia, which is higher than the number of SWPs as 2. This might be due to the fact the small amount of sequence submitted from South Asia in current time.

### Mutations on S protein increase binding capacity of ACE2 and decrease affinity of antibody

The S protein of the SARS-CoV-2 is the major target for the humoral immunity and the receptor-binding domain (RBD) of S protein could mediate the attachment of viruses to the surface receptors in host cell^25-27^. Thus, under the selective pressure of both sides, the S protein was mutated frequently. Here, we evaluate the mutations on S protein, specifically on the ACE2 binding domain and the epitopes for antibody. The 3-D structures of three mutants with mutation in ACE2 binding domain including R408I, N439K, G476S (Figure XX) and four mutants with mutation in epitope regions including K378R, H519P, H519Q, R408I (**Figure 2d**) were constructed by homology modeling^28^. Further, the binding affinities between corresponding mutants and ACE2^29^ or antibody CR3022^30^ were calculated through molecular docking^31^. The RBD region of SARS-CoV-2 isolate Wuhan-Hu-1(MN908947.3) was selected as reference for control.

As been illustrated in **Figure 2d**, the mutants of R408I, N439K and G476S could promote the binding capacity between RBD region and ACE2 compared with reference. Among them, arginine (R) and lysine (K) are alkaline amino acids, the mutations of R to isoleucine (I) and asparagine (N) to K involving significant property changes. Further investigation illustrated the potential binding site for N439K is the acidic amino acids glutamicacid in 329E, which could form an ionic bond to increase the binding capacity between RBD and ACE2 (**Figure 2d**). Moreover, the potential binding site for R408I is the alkaline amino acid of histidine (H), the mutations of R408I might reduce the repulsive force between two alkaline amino acids of H and R (**Figure 2d**).

On the contrary, the mutations of K378R, H519P, H519Q, R408I could decrease the binding capacity of antibody CR3022. Among them, only K378R didn’t involving huge property changes since K and R are both alkaline amino acids. Other three are both alkaline amino acids mutated to uncharged residues. It could be observed that 408R could form an ionic bond with the asparticacid (D) in 54D on the H chain. After R mutated to I, the ionic bond will be broken to reduce the binding capacity of the antibody. Above results indicated that the mutations on the RBD of S protein might enhance the ability of virus to target the ACE2 receptor and further promote the capacity of virus to invade host cell. Moreover, the mutations on RBD hold the potential to decrease the binding ability of antibodies and lead to immune escape of the virus.

## Discussion

At present, the global pandemic of SARS-CoV-2 immeasurably impact the human health and economic development worldwide. Thus, revealing the evolutionary pattern would be extremely helpful for the disease prevention and vaccine design. In this article, we comprehensively analysis the mutation pattern based on the largest available dataset of mutations to reveal the evolution direction and potential drivers of SARS-CoV-2. Results showed that the mutations could cause more WSPs than SWPs on the whole genome. However, for cross-reactive peptide between seasonal HCoVs and SARS-CoV-2, the number of SWPs are significantly higher than WSPs on ORF1ab, indicating the potential natural selection of SARS-CoV-2 by pre-existing cross-reactive T cell immunity. Further, the analysis on S protein illustrated that the evolution of SARS-CoV-2 was more likely to be more infective by increase the binding affinity of ACE2 and escape the surveillance of the immune system by reduce the binding affinity of the corresponding antibody.

Since Dec, 2019, the SARS-CoV-2 has spread to over 200 countries or regions in six months, and maintained the continuous transmission so far. The long terms community transmission in different regions and races will make SARS-CoV-2 under different the types of immune pressure. As a typical RNA virus, those selective pressure will drove the frequent mutations of SARS-CoV-2, moreover, according to our results, this evolution may have certain directionality towards immune escape.

Currently, nations such as United States, United Kingdom, and China are suffering from the epidemic and are the high incidence areas of SARS-CoV-2. It can be observed that the long-term and large-scale community transmission in these countries makes the existential time of weak binding peptides longer than strong binding peptides. It is suggested that the SARS-CoV-2 in those countries may have the evolutionary direction of reducing the immunogenicity of T-cell epitopes. Furthermore, we systematically analyzed the mutation of CREs between SARS-CoV-2 and seasonal HCoVs. The results showed that although the number of SWPs and WSPs in the mutations of whole genome peptides are not significantly different from each other, however, SWPs in the mutations of CREs had an overwhelming number over WSPs in tens or even hundreds of times. Since adults will be infected with seasonal HCoVs every two to three years^7^, seasonal HCoVs have already been spread with in large-scale of population in a long-term. Therefore, the mutations of CREs in SARS-CoV-2 will evolved towards the direction of immune escape.

Finally, we analyzed the target protein Spike of the B-cell immunity and found that the S protein of SARS-CoV-2 can not only enhance the binding affinity with human receptor ACE2 by form a new ionic bond through mutation such as N439K, but also reduce the binding affinity of antibody CR3022 by destroying the ionic bond through mutation R408I. This result suggests that S protein has an evolutionary trend to increase infectious ability and escape immune monitoring under B-cell immune pressure.

According to a latest research of the SARS-CoV-2 transmission dynamics^18^, SARS-CoV-2 is more likely to spread much longer than we expected. This research points out that if the immune response for SARS-CoV-2 infections are long-term or even life-time effective, the epidemic may end in year 2021. However, if the immune memory is short-term effective, such as 40 weeks to 104 weeks, the risk of annual or periodic eruption in every two or three years is existed, like the seasonal HCoVs. Consistently, our research indicated that rapid mutations of SARS-CoV-2 may trend to escape from the monitoring and recognition of the immune system. This means that even if the effective vaccine could be developed for current circulating SARS-CoV-2, the rapid, immune escape trend mutations will cause the ineffective of vaccine in a short time. Thus, we may expect the vaccine development become a cyclical work for SARS-CoV-2 in the future, just like the case of influenza virus. In this circumstance, monitoring the mutations and antigenic evolution of SARS-CoV-2 timely will be a very important task.

## METHODS

### Data source

The reference amino acid sequences of SARS-CoV-2 (MN908947.3) and four human susceptible seasonal CoVs including HCoV-229E (NC_002645.1), HCoV-HKU1 (NC_006577.2), HCoV-NL63 (NC_005831.2), and HCoV-OC43 (NC_006213.1) were derived from the National Center for Biotechnology Information (NCBI)^32^.

The mutant information (GFF3 files) of 10 structure and non-structure proteins including spike protein, envelope protein, nucleocapsid protein, membrane protein, orf1ab protein, ORF3a, OFR6, ORF7a, ORF8 and ORF10 were derived from the National Genomics Data Center^33^ till May, 20^th^, 2020. For mutation analysis of the whole protein sequence, we extended the indel positions to the reference sequence to obtain a more precise mutation frequency for each amino acid point. For ease of k-mer analysis, all non-mismatch mutations are excluded, which left 4,420 mutation sites for downstream analysis.

Frequency of alleles in different countries and regions were obtained from Allele Frequency Net Database (AFND)^34^. For each country and region, only those alleles with frequency over 0.05 and population over 25 were retained.

### Peptide preparation

#### Mutant peptide preparation

For each mutation site *i* compared with reference sequence Wuhan-Hu-1(MN908947.3), the sequence from site *i* − 15 to *i* + 15 was derived from the corresponding protein sequence of the mutants. Further, it will be divided into peptides with the sliding windows of 8-mer to 15-mer and searching step of 1-mer. Among all peptides, those contain the mutation site *i* was retained as the mutant epitopes.

#### Cross-reactive epitope preparation

Each of the mutant peptide of SARS-CoV-2 was compared against all four seasonal CoVs using in-house python script. The cross-reactive epitopes were defined as those peptides that are identical to the k-mer sequence in at least one of the seasonal CoVs.

### Prediction of HLA binding affinity

Both the HLA-I and HLA-II binding affinities between HLA alleles and k-mer peptides were predicted by the T Cell Epitope Prediction Tools (standalone version 2.22.3) with IEDB (Immune Epitope Database) recommended methods^21,20^. For HLA I alleles, 8-14 mer peptides were analyzed and were considered strong binding peptides with a cutoff ≤ 0.01^35^. For HLA II alleles, only 15 mer peptides were analyzed and a consensus percentile rank of the top 10% was considered strong binding peptide^36^. The allele-peptide pairs with prediction score lager than the cutoff was considered weak binding peptides. The total number of strong or weak binders were determined by the combination of strains and alleles.

## Author Contributions

T.Y.Q., Q.B.L, and C.D.Z. design the experiments. C.D.Z. and X.X.J collected the data and perform the analysis. X.X.J. and C.X.Y. organize the results. T.Y.Q., and Q.B.L. co-supervise the whole project. All of them contributing to the writing of the manuscript.

## Declaration of interests

The authors declare no competing interests.

## Acknowledgements

This work is supported in part by grants from the National Natural Science Foundation of China (31900483), the Shanghai Sailing program (19YF1441100) and the Shanghai Public Health Clinical Center (KY-GW-2020-07), the National Key Research and Development Program of China (SQ2018YFA090045-01).

## Notes

### Competing Interest Statement

The authors have declared no competing interest.

