## Supplementary Files for "Antigenic evolution on global scale reveals potential natural selection of SARS-CoV-2 by pre-existing cross-reactive T cell immunity"

**Supplementary Table**

**Supplementary Table 1. Mutation Frequency on the whole genome of SARS-CoV-2^*^.**

^*^Each line illustrated the protein name, mutation positions, mutation counts, mutation frequency and whether is it in the conserved regions.

**Supplementary Table 2. Top ranking mismatch mutations on the whole genome of SARS-CoV-2^*^.**

^*^Each line illustrated the protein name, mutation positions, number of mutation types, mutation counts and counts of different mutation types.

**Supplementary Table 3. Top ranking mutations on the circulating strains in different regions^*^.**

^*^Each lines illustrated the protein name, mutation positions, mutation types, circulating countries, mutation counts and circulating continent.

**Supplementary Table 4. Numbers of four types of HLA-I PTEs before and after the mutations**^*^**.**

| **All** | | **Protein** | | | |
| --- | --- | --- | --- | --- | --- |
| strong→weak： | 896351 | **orf1ab** |  | **orf10** |  |
| weak→strong： | 1339795 | strong→weak： | 628847 | strong→weak： | 4618 |
| strong→strong： | 2176811 | weak→strong： | 1007040 | weak→strong： | 2527 |
| weak→weak： | 361873335 | strong→strong： | 1610993 | strong→strong： | 8955 |
|  |  | weak→weak： | 183483484 | weak→weak： | 529616 |
|  |  | **orf3a** |  | **E protein** |  |
|  |  | strong→weak： | 81690 | strong→weak： | 623 |
|  |  | weak→strong： | 177343 | weak→strong： | 2145 |
|  |  | strong→strong： | 353207 | strong→strong： | 1849 |
|  |  | weak→weak： | 40811273 | weak→weak： | 402090 |
|  |  | **orf6** |  | **S protein** |  |
|  |  | strong→weak： | 1311 | strong→weak： | 110643 |
|  |  | weak→strong： | 1628 | weak→strong： | 54221 |
|  |  | strong→strong： | 2729 | strong→strong： | 74905 |
|  |  | weak→weak： | 624377 | weak→weak： | 70248776 |
|  |  | **orf7a** |  | **M protein** |  |
|  |  | strong→weak： | 4349 | strong→weak： | 7837 |
|  |  | weak→strong： | 5812 | weak→strong： | 13688 |
|  |  | strong→strong： | 11058 | strong→strong： | 32425 |
|  |  | weak→weak： | 1580228 | weak→weak： | 2695335 |
|  |  | **orf8** |  | **N protein** |  |
|  |  | strong→weak： | 37905 | strong→weak： | 18528 |
|  |  | weak→strong： | 18198 | weak→strong： | 57193 |
|  |  | strong→strong： | 38049 | strong→strong： | 42641 |
|  |  | weak→weak： | 18461323 | weak→weak： | 43036833 |

*Here, we defined four types of predicted HLA-I PTEs. The predicted binding affinity was defined as strong or weak according to defaulted threshold before or after the mutations. For example, strong🡪weak refers to the peptide predicted as strong binding before the mutation and alter to weak binding after the mutation.

**Supplementary Table 5. Numbers of four types of HLA-II PTEs before and after the mutations**^*^**.**

| **All** | | **protein** | | | |
| --- | --- | --- | --- | --- | --- |
| strong→weak： | 1901868 | **orf1ab** |  | **orf10** |  |
| weak→strong： | 6834805 | strong→weak： | 951691 | strong→weak： | 8272 |
| strong→strong： | 11687013 | weak→strong： | 5285717 | weak→strong： | 13954 |
| weak→weak： | 131568872 | strong→strong： | 8947725 | strong→strong： | 35716 |
|  |  | weak→weak： | 60722092 | weak→weak： | 162715 |
|  |  | **orf3a** |  | **E protein** |  |
|  |  | strong→weak： | 298075 | strong→weak： | 1810 |
|  |  | weak→strong： | 962867 | weak→strong： | 7861 |
|  |  | strong→strong： | 1723479 | strong→strong： | 13696 |
|  |  | weak→weak： | 13098978 | weak→weak： | 131269 |
|  |  | **orf6** |  | **S protein** |  |
|  |  | strong→weak： | 8854 | strong→weak： | 462542 |
|  |  | weak→strong： | 6986 | weak→strong： | 287644 |
|  |  | strong→strong： | 61716 | strong→strong： | 567467 |
|  |  | weak→weak： | 177087 | weak→weak： | 28835652 |
|  |  | **orf7a** |  | **M protein** |  |
|  |  | strong→weak： | 8524 | strong→weak： | 12605 |
|  |  | weak→strong： | 25125 | weak→strong： | 98848 |
|  |  | strong→strong： | 46353 | strong→strong： | 140104 |
|  |  | weak→weak： | 581295 | weak→weak： | 1015623 |
|  |  | **orf8** |  | **N protein** |  |
|  |  | strong→weak： | 27215 | strong→weak： | 122280 |
|  |  | weak→strong： | 71401 | weak→strong： | 74402 |
|  |  | strong→strong： | 44167 | strong→strong： | 106590 |
|  |  | weak→weak： | 6527958 | weak→weak： | 20316203 |

*Here, we defined four types of predicted HLA-II PTEs. The predicted binding affinity was defined as strong or weak according to defaulted threshold before or after the mutations. For example, strong🡪weak refers to the peptide predicted as strong binding before the mutation and alter to weak binding after the mutation.

**Supplementary Table 6. Numbers of SWPs and WSPs cross-reactive peptide between seasonal HCoVs and SARS-CoV-2**^*^**.**

| **HLA-I** | | **HLA-II** |
| --- | --- | --- |
| **area: North America** | **area: South Asia** | **area: North America** |
| protein orf1ab | protein orf1ab | protein orf1ab |
| weak→strong： 1733 | weak→strong： 61 | weak→strong： 160 |
| strong→weak： 34800 | strong→weak： 135 | strong→weak： 1024 |
| **area: North America** | **area: East Asia** | **area: Europe** |
| protein n | protein orf1ab | protein orf1ab |
| weak→strong： 4 | weak→strong： 85 | weak→strong： 357 |
| strong→weak： 0 | strong→weak： 148 | strong→weak： 870 |
| **area: Europe** | **area: Southeast Asia** | **area: East Asia** |
| protein orf1ab | protein orf1ab | protein orf1ab |
| weak→strong： 732 | weak→strong： 1 | weak→strong： 16 |
| strong→weak： 1282 | strong→weak： 4 | strong→weak： 310 |
|  | **area: Oceania** | **area: South Asia** |
|  | protein orf1ab | protein orf1ab |
|  | strong→weak： 7 | weak→strong： 13 |
|  |  | strong→weak： 2 |

*Here, we provide the available number of weak to strong peptides (WSPs) and strong to weak peptide (SWPs).

**Supplementary Table 7. SWPs of CRPs between seasonal HCoVs and SARS-CoV-2**^*^**.**

*Here, we presented all SWPs with counting number of 10 in our dataset. Each line represents the protein name, peptides on the reference protein, peptides on the mutants, mutations, start and end positions on the whole genome.
